# Cryo-EM structures of human TMEM120A and TMEM120B

**DOI:** 10.1101/2021.06.27.450060

**Authors:** Meng Ke, Yue Yu, Changjian Zhao, Shirong Lai, Qiang Su, Weidan Yuan, Lina Yang, Dong Deng, Kun Wu, Weizheng Zeng, Jia Geng, Jianping Wu, Zhen Yan

## Abstract

TMEM120A (Transmembrane protein 120A) was recently identified as a mechanical pain sensing ion channel named as TACAN, while its homologue TMEM120B has no mechanosensing property^1^. Here, we report the cryo-EM structures of both human TMEM120A and TMEM120B. The two structures share the same dimeric assembly, mediated by extensive interactions through the transmembrane domain (TMD) and the N-terminal coiled coil domain (CCD). However, the nearly identical structures cannot provide clues for the difference in mechanosensing between TMEM120A and TMEM120B. Although TMEM120A could mediate conducting currents in a bilayer system, it does not mediate mechanical-induced currents in a heterologous expression system, suggesting TMEM120A is unlikely a mechanosensing channel. Instead, the TMDs of TMEM120A and TMEM120B resemble the structure of a fatty acid elongase, ELOVL7, indicating their potential role of an enzyme in lipid metabolism.

## Introduction

Pain is a protective signal of impending danger, which is essential for our interaction with the environment^2^. The identification of related mechanosensing ion channel (MSC) is fundamentally important for understanding the mechanism of mechanical pain sensing and analgesic drug discovery. A recent study identified TMEM120A, also named as TACAN, as a MSC that contributes to mechanosensitive currents in nociceptors^1^. TMEM120A is novel and shares no sequence homology with any reported MSCs or other channels. Despite sequence similarity, no mechanically evoked current was detected for heterologously expressed TMEM120B, a homologue of TMEM120A^1^. TMEM120A/B were originally identified as a nuclear membrane localized protein that plays an important role in adipocyte differentiation^3,4^. A recent study shows that adipocyte-specific knockout of TMEM120A leads to latent lipodystrophy^5^. The molecular basis underlying the physiological functions of TMEM120A is elusive. Here, we present the cryo-electron microscopy (cryo-EM) structures of both human TMEM120A and TMEM120B at an overall resolution of 3.7 Å and 4.0 Å, respectively. Unexpectedly, the almost identical structures between TMEM120A and TMEM120B, and our electrophysiological characterizations do not support the role of TMEM120A as an MSC. Instead, the TMD of TMEM120A/B share similar fold with ELOVL7, a member of the ELOVL family elongase catalyzing the first step in the fatty acid elongation cycle^6^. These clues suggest TMEM120A/B are likely enzymes involved in lipid metabolism, although its detailed role remains to be investigated.

## Results

### Human TMEM120A and TMEM120B are homodimers

We sought to a comparative structural investigation between TMEM120A and TMEM120B, trying to uncover their potential difference in mechanosensing property. Details about the sample preparation, data collection, and processing can be found in Supplementary figures and Methods (Figs. S1-3). The cryo-EM maps of TMEM120A and TMEM120B reveal a shared homo-dimeric assembly, which is about 80 Å in height and 85 Å in width (Fig. 1a). The two structures are nearly identical with a root-mean-square distance (r.m.s.d.) of only 0.82 Å over 293 Cα atoms when superimposed with each other (Fig. 1b).

**Fig. 1.**
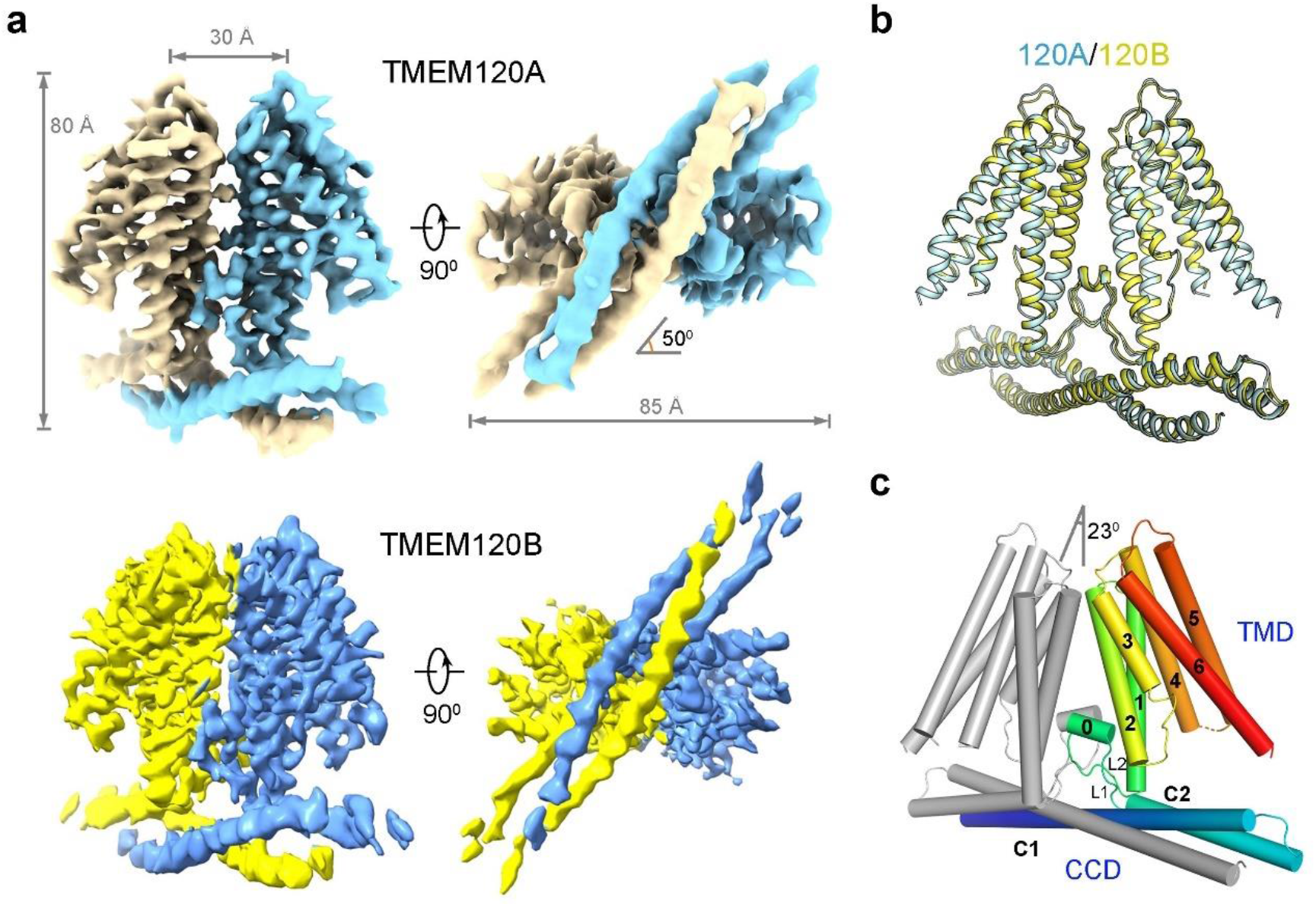
Overall structures of human TMEM120A and TMEM120B. **a**, Cryo-EM maps of human TMEM120A and TMEM120B shown in two perpendicular views. Densities for the detergent micelle were omitted for visual clarity. The maps were contoured at level 0.7 and 0.6 for TMEM120A and TMEM120B in ChimeraX^18^, respectively. **b**, TMEM120A and TMEM120B share nearly identical overall structure. TMEM120A and TMEM120B are colored cyan and yellow, respectively. **c**, Structural topology of TMEM120A. One protomer is colored gray and the other is colored rainbow with the N- and C-terminus in blue and red, respectively. TMD: transmembrane domain; CCD: coiled coil domain. Structure figures were prepared in PyMOL^19^.

We select TMEM120A for detailed analysis unless otherwise stated. Both the N-terminus and the C-terminus of each protomer are located on the intracellular side. The TMD of each protomer contains six transmembrane helices, designated H1-H6, enclosing a large solvent accessible cytosol-facing cavity. The N-terminal cytosolic region from both protomer together form the CCD, with each protomer contributing two horizontal helices C1 and C2 (Fig. 1c, Fig. S4). In addition, a short amphipathic helix H0 parallel to the inner membrane was identified between C2 and H1, flanked by two intracellular linkers L1 and L2 (Fig. 1c). When viewed from the cytosolic side, long axes of CCD and TMD form an angle of approximately 50° (Fig. 1a).

The dimeric assembly is stabilized by extensive interactions in both CCD and TMD (Fig. 2a). The C1 helices from the two protomers form an anti-parallel coiled coil through multiple types of interactions, including hydrogen bonds, cation-π interactions and van der Waals contacts. At the two ends of the coiled coil, the short C2 helix jointly to form two 3-helix bundles (Fig. 1c, 2a, b). The dimer interface in TMD, which is along the C2 symmetry axis, is mostly mediated by van der Waals contacts. Near the extracellular side, Val159 and Phe166 on H2, Leu208 on H3, and Trp305 on H6 from the two protomers contribute to the interface interaction. On the interface near the intracellular side, three layers of hydrophobic residues from L2, H0 and H2 interlock with each other (Fig. 2c). In addition, Arg178 on H2 of one protomer forms hydrogen bonds with the carbonyl oxygens of Leu111 and Val112 on H0’ of the other protomer. These specific interactions stabilize H0, which in turn buttress dimerization (Fig. 2b).

**Fig. 2.**
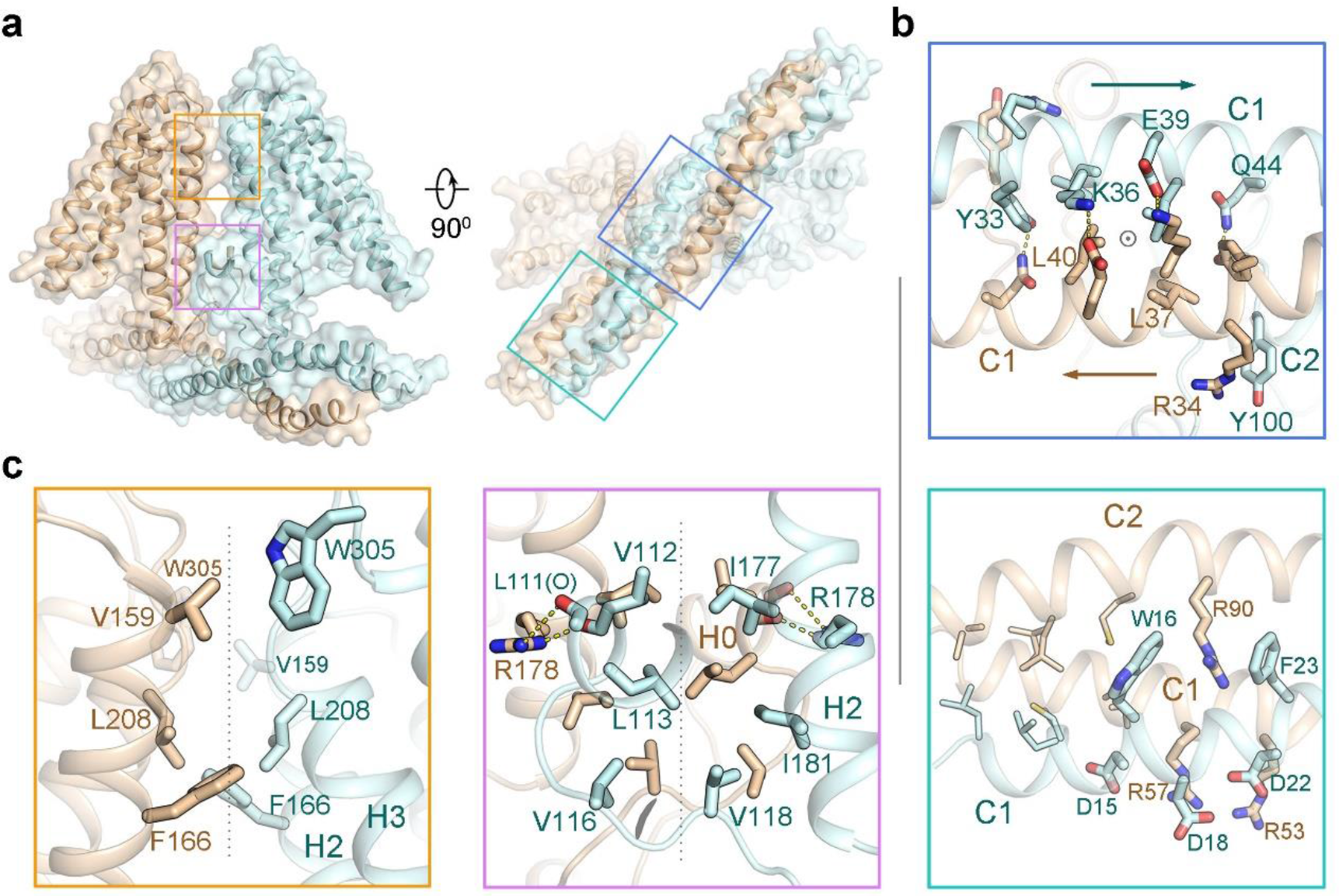
Homo-dimeric assembly of TMEM120A. **a**, The two TMEM120A protomers interact extensively through both TMD and CCD. The structure of TMEM120A, presented as cartoon under semi-transparent surface, is shown in two perpendicular views. The dimer interfaces that are highlighted in the boxes are illustrated in detail in panels b and c with corresponding color borders. **b**, The dimeric interface of TMD. The interfaces are along the C2 symmetry axis, which is indicated by the gray dashed lines. **c**, The dimeric interface of CCD. The C2 symmetry axis is indicated by the circle in the upper panel.

Most of the residues in the dimer interface are highly conserved between TMEM120A and TMEM120B (Fig. S5). The CCD is the less conserved region, but the overall structures are similar. From the structural perspective, there is hardly any hint for the difference between TMEM120A and TMEM120B in sensing mechanical force.

### Electrophysiological characterizations of human TMEM120A

We also examined the electrophysiological property of TMEM120A in both bilayer system and heterologous expression system. We purified the full length human TMEM120A and TMEM120B and reconstituted them into a planar lipid bilayer for conducting current recording. A discrete current increase, measured at the voltage of 10 mV, was observed after reconstitution of purified TMEM120A (Fig. S6a). TMEM120A showed an almost linear current-voltage (I-V) relationship between −50 mV and +50 mV, which is consistent with the reported electrophysiological behavior in bilayer^1^ (Fig. S6b, c). We also examined TMEM120B using the same system, and the current it mediated was slightly smaller than that by TMEM120A at the same voltage level (Fig. S6c).

We tested the mechano-sensing property of TMEM120A by two mechanical force modes in a heterologous expression system, in which the plasmids of TMEM120A were transiently transfected into HEK293T or HEK293T-P1KO cells. Under the whole-cell poking mode, we applied steps of mechanical stimuli on the measured cells by a fire-polished glass pipette with 200 milliseconds duration. Cells transfected with mouse Piezo1 showed significantly increased mechanically evoked currents compared with the empty vector transfected cells (Fig. 3a). By contrast, no significant difference was detected between cells transfected with TEME120A/B and the empty vector-transfected cells (Fig. 3a, b). We also measured the stretch-induced currents by applying negative pressures under the cell-attached stretch mode. The positive control cells transfected with human Piezo1 mediated robust stretch-induced currents and the amplitude of the currents were improved with the increasing stretch pressure (Fig. 3c, d). Out of ten measured cells transfected with TMEM120A, six showed no obvious stretch-induced currents, similar to the negative control cells transfected with vector. The other four cells displayed currents in ~ 5 picoampere level, which is likely to be leaky currents for two reasons: First, the amplitude of the current is not proportional to the stretch pressures. Second, the currents exist even after the pressure is relieved. Therefore, based on our experimental results, TMEM120A could mediate conducting currents in a bilayer system, but does not behave as an MSC in heterologous system.

**Fig. 3.**
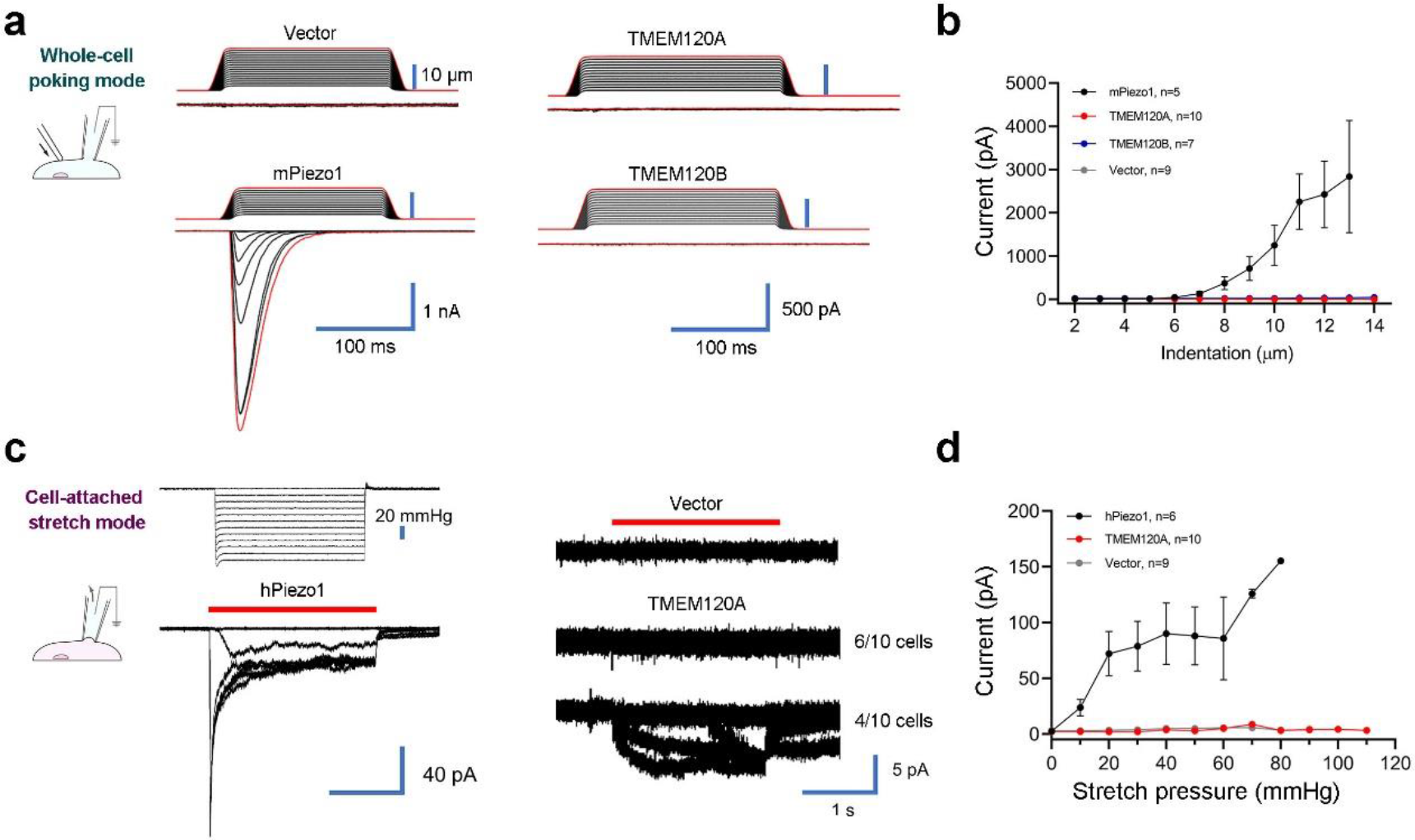
TMEM120A does not produce mechanically induced currents in heterologous system. **a**, Representative traces of poking-evoked whole cell currents from HEK293T-P1KO cells transfected with mPiezo1 and TMEM120A, or from HEK293T cells transfected with TMEM120B, and vector. **b**, Statistic plot of the poking-evoked currents along the increase of indentation depth. **c**, Representative traces of stretch induced currents in a cell-attached mode from HEK293T-P1KO cells transfected with mPiezo1, TMEM120A, and vector. The stretch durations are indicated by red bars. **d**, Statistic plot of the stretch-induced currents along the increase of negative pressure.

### TMEM120A shared a common fold with ELOVL7

A search of TMEM120A protomer in the DALI^7^ server identified a structurally homologous protein ELOVL7^6^, an endoplasmic reticulum embedded fatty acid elongase involves in the <u>elo</u>ngation of <u>v</u>ery long chain fatty acid. ELOVL7 catalyzes the condensation reaction step between an acyl-CoA and malonyl-CoA to form a 3-keto acyl-CoA, followed by three other steps to yield a product acyl-CoA with two extra carbons. ELOVL7 contains seven TM helices, TM1-TM7. Structural comparison between TMEM120A and ELOVL7 reveals similar fold in the TMD. The H1-H6 of TEME120A can superimpose with TM2-TM7 of ELOVL7, with an r.m.s.d. of about 2.8 Å over 193 aligned Cα atoms (Fig. 4a). Notably, two histidine residues His150 and His151 of the HxxHH motif in the catalytic site of ELOVL7 are also conserved in TMEM120A (Fig. 4b). In addition, like ELOVL7, the cytosol-facing cavity of TMEM120A is surrounded by positively charged residues. An extra density was observed in the cavity, the size and shape of which is reminiscent of the 3’-phoshoadenosine group of acyl-CoA. TMEM120A is unlikely to have the same function as ELOVL7, as another key residue His181 in the catalytic site of ELOVL7 is not conserved in TMEM120A (Fig. 4b). Nevertheless, the similarity of the overall fold, cytosol-facing positively charge pocket, and a potential bound molecule in the cavity imply TMEM120A may be an enzyme in lipid metabolism that catalyze a different reaction from ELOVL7.

**Fig. 4.**
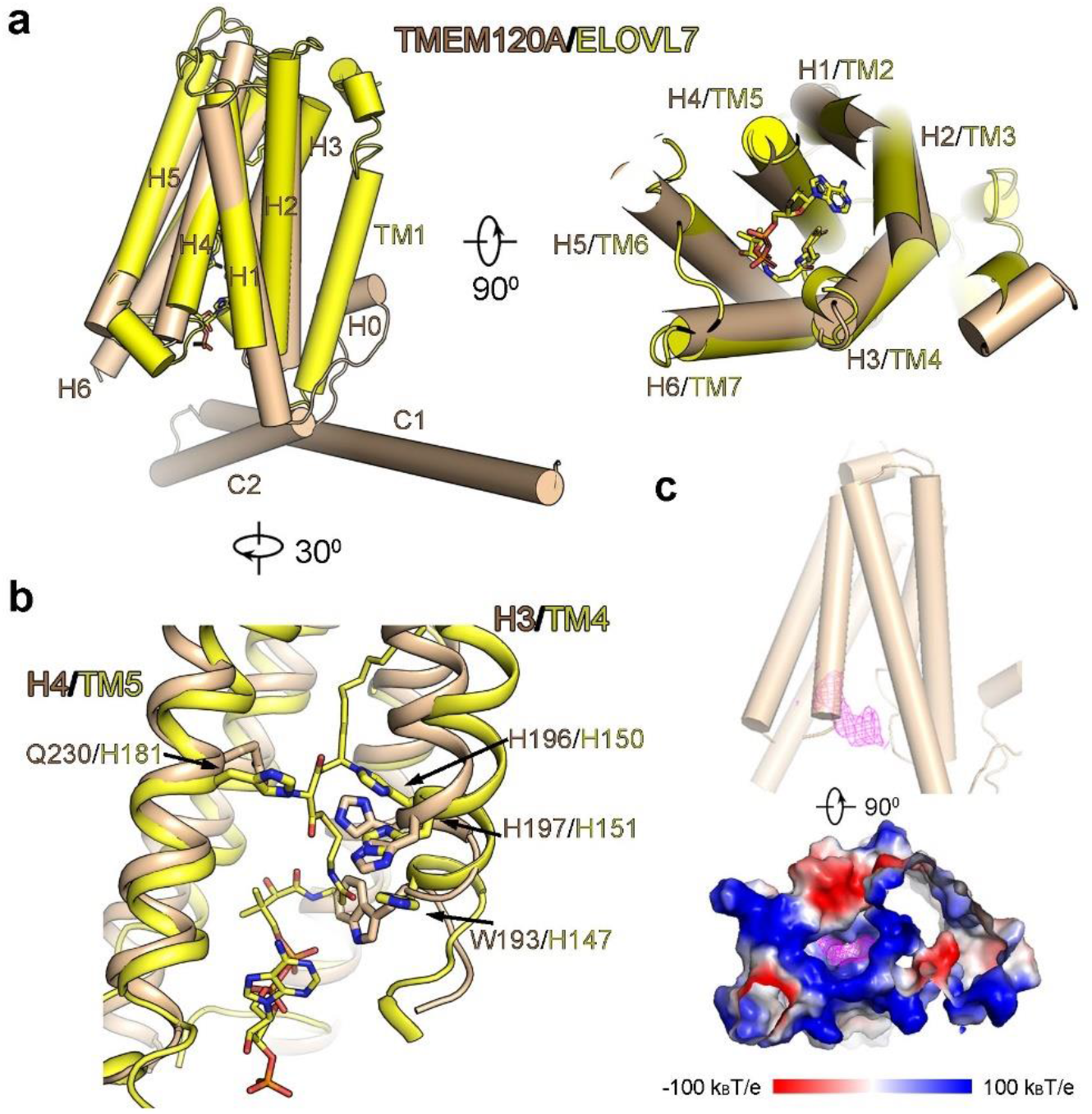
The structure of TMEM120A TMD resembles that of ELOVL7 elongase. **a**, Structural comparison between TMEM120A and ELOVL7 elongase (PDB: 6Y7F), shown as cylindrical helices in two perpendicular views. The 3-keto acyl-CoA bound in ELOVL7 is shown as sticks. **b**, Close-up view of the superimposition near the HxxHH motif in the catalytic site of ELOVL7. **c**, A stretch of electron density is identified in the intracellular pocket of TMEM120A, which is surrounded by positively charged residues. The density is contoured at 4σ in PyMOL.

## Discussion

In this study, we examined the electrophysiological property in both bilayer and heterologous expression systems and performed a comparative structural investigation on TMEM120A and TMEM120B. The two nearly identical structures do not provide any hint for their difference in the ability of mechanosensing as suggested by previous study^1^. The overall structure of TMEM120A revealed in our study is not similar to any previously reported MSC structures. The ion permeation path of most MSCs, including MscL and MscS^8^ from bacteria, NOMPC^9^ from fly, and TRAAK^10^ and Piezo^11,12^ from mammal, are along the symmetric axis. The dimeric TMEM120A is reminiscent of OSCAs^13–16^ from plant, whose ion permeation path are along each protomer. Although TMEM120A shows the ability to permeate ions as measured in bilayer system, our results indicate TMEM120A does not response to poking or stretch mechanical stimuli in heterologous expression system. We even cannot conclude TMEM120A is an ion channel as conducting currents were also measured for several non-channel proteins in bilayer system^17^. The structural similarity between TMEM120A and ELOVL7, and the extra density bound in the cytosol-facing cavity suggest that TMEM120A may function as an enzyme in lipid metabolism^3^. However, the exact function of TMEM120A remains to be further investigated.

## Acknowledgements

We would like to thank the Cryo-EM Facility and the HPC Center of Westlake University for providing cryo-EM and computation support. This work was supported by Westlake Education Foundation to Z.Y.. Atomic coordinates and corresponding EM maps of the human TMEM120A (PDB: 7CXR; EMDB: EMD-30495) and TMEM120B (PDB: 7F73; EMDB: EMD-31484) have been deposited in the Protein Data Bank (http://www.rcsb.org) and the Electron Microscopy Data Bank (https://www.ebi.ac.uk/pdbe/emdb/), respectively.

