## supplementary information for "Cryo-EM structures of human TMEM120A and TMEM120B"

**Materials and Methods**

**Protein sample preparation**

The codon optimized gene of human TMEM120A and TMEM120B were synthesized and subcloned into pCAG vector with a N-terminal FLAG and mCherry tag. HEK 293F cells (Invitrogen) were transiently transfected with the plasmids and harvested after 60 hours. For each batch of purification, 2-liter cells were pelleted and resuspended in buffer containing 25 mM Tris-HCl (pH 8.0), 300 mM NaCl, 0.5% (w/v) Lauryl Maltose Neopentyl Glycol (LMNG, Anatrace), 0.06% (w/v) cholesteryl hemisuccinate Tris salt (CHS, Anatrace) and protease inhibitor cocktail including 2 mM phenylmethylsulfonyl fluoride (PMSF), 1.3 μg/ml aprotinin, 0.7 μg/ml pepstatin, and 5 μg/ml leupeptin. The mixture was incubated at 4°C for 2 hours on a rotating shaker. The insoluble fraction was precipitated by ultracentrifugation at 255,700 g for 1 h, and the supernatant was applied to anti-Flag G1 affinity resin (GenScript) by gravity. The resin was washed sequentially to remove unbound proteins with wash buffer containing 25 mM Tris-HCl (pH 8.0), 400 mM NaCl, 0.015% (w/v) glyco diosgenin (GDN, Anatrace), and protease inhibitor cocktail. The target protein was eluted by the elution buffer containing 25 mM Tris-HCl (pH 8.0), 400 mM NaCl, 0.015% (w/v) GDN, and protease inhibitor cocktail plus 250 μg/ml FLAG peptide (GenScript). The eluent was then concentrated using a 30-kDa cut-off Centricon (Millipore) and further purified by size exclusion chromatography (Superose 6 Increase 10/300 GL column, GE Healthcare) in 25 mM Tris-HCl (pH 8.0), 150 mM NaCl, 0.015% (w/v) GDN, and protease inhibitor cocktail. Peak fractions containing the target protein were collected and concentrated to about 6.5 mg/mL for cryo-EM analysis. The wild type and mutant proteins for electrophysiological experiments in lipid bilayer were prepared in a similar procedure except for the detergent used in the gel filtration was replaced by 0.05% Lauryl dimethylamine-N-Oxide (LDAO, Anatrace) and 0.01% CHS.

**Bilayer recording**

Electrophysiological recordings of wild type TMEM120A, TMEM120B, and TMEM120A mutants were performed on the planar bilayer lipid membrane formed in a cup (1 mL) and a chamber (1 mL) (Warner instrument), with the buffer conditions of *cis*-: 500 mM NaCl, 10 mM HEPES, pH 7.5 and *trans*-: 100 mM NaCl, 10 mM HEPES, pH 7.5. *E.coli* Polar Lipid Extract (AVANTI) was dissolved in decane with a final concentration of 25 mg/mL. The aperture (150 μm) of the cup was precoated with 0.8 μL lipid solution. After the bilayer lipid membrane was formed, 1 μL of purified protein (about 6 mg/mL) was added into the *cis-* chamber. Current trace was recorded by HEKA EPC 10 USB (HEKA) and Axonpatch 200B (Axon Instruments) with 10 kHz sampling frequency.

**Cells and transfections**

HEK293T-P1KO cells used for whole-cell electrophysiology is a gift from Dr. Bailong Xiao and HEK293T-P1KO cells used for cell-attached electrophysiology were generated using CRISPR-Cas9 technique as previously described^1^^,^^2^. The HEK293T or HEK293T-P1KO cells were cultured in Dulbecco's Modified Eagle Medium (DMEM, BI) containing 4.5 mg/mL glucose, 10% fetal bovine serum (FBS, BI) and co-transfected with the expression plasmids or vector when cell confluency reached 70% using lipofectamine 2000 (Invitrogen). After incubation at 37 ^o^C under 5% CO_2_ for 24 hours, the cells were treated with 0.05% trypsin (BI) and put on poly-D-lysine (Sigma-Aldrich) coated 12 mm cover slips (Assistent) for electrophysiological characterization. No further authentication was performed for the commercially available cell line. Mycoplasma contamination was not tested. All plasmids were transfected at a concentration of 600 ng/mL. Cell were recorded from 24 to 48 hr after transfection.

**Whole cell electrophysiology**

The whole-cell currents were recorded using an EPC-10 amplifier with Patchmaster 2.90.4 software (HEKA Elektronik) and glass micropipettes (2-4 MΩ, Sutter Instrument) made by P-97 pipette puller (Sutter Instrument). The electrodes were filled with the internal solution composed of 140 mM KCl, 8 mM NaCl, 10 mM HEPES, 4 mM MgATP and 0.5 mM Na_2_GTP, pH 7.2 with KOH, and the extracellular solution was composed of 130 mM NaCl, 20 mM KCl, 1 mM CaCl_2_, 2 mM MgCl_2_, 5 mM D-Glucose, 10 mM HEPES, pH 7.2 with NaOH. All experiments were performed at room temperature. Currents were sampled at 20 kHz and filtered at 2 kHz. Leak currents before mechanical stimulations were subtracted offline from the current traces. The data were analyzed using Fitmaster 2.90.4 (HEKA Elektronik) and Prism 8.2 (GraphPad Software).

Mechanical stimulation utilized a fire-polished glass pipette (tip diameter 3–4 mm) positioned at an angel of 80° relative to the cell being recorded as described^3^. The probe was driven by Patchmaster-controlled piezoelectric crystal microstage (E625 LVPZT Controller/Amplifier; Physik Instrument). The probe velocity was 1 μm/ms during the upward and downward movement, and the stimulus was kept constant for 200 ms. A series of mechanical steps in 1μm increments was applied every 10 s and currents were recorded at a holding potential of -70 mV.

**Cell-attached electrophysiology**

For cell-attached patch clamp recordings, external solution used to zero the membrane potential consisted of 140 mM KCl, 1 mM MgCl_2_, 10 mM glucose and 10 mM HEPES (pH 7.3 with KOH). Recording pipettes were of 1 – 3 MΩ resistance when filled with standard solution composed of 130 mM NaCl, 5 mM KCl, 1 mM CaCl_2_, 1 mM MgCl_2_, 10 mM TEA-Cl and 10 mM HEPES (pH 7.3 with NaOH). Stretch-activated currents were recorded in the cell-attached patch clamp configuration using Axopatch 200B amplifier (Axon Instruments). Membrane patches were stimulated with 2 s negative pressure pulses through the recording electrode using Clampex controlled pressure clamp HSPC-1 device (ALA-scientific), with inter-sweep duration of 30s. Currents were sampled at 20 kHz and filtered at 2 kHz or 10 kHz. Leak currents before mechanical stimulations were subtracted off-line from the current traces. All experiments were done at room temperature. All data was analyzed in Clampfit 11.2 and data was plotted using GraphPad Prism.

**Cryo-EM sample preparation and data collection**

0.1% Fos-Choline-8, Fluorinated (Anatrace) was added into the protein sample before grid preparation. Aliquots (3.5 μL) of purified TMEM120A or TMEM120B with concentration about 6.5 mg/ml were loaded onto glow-discharged holey carbon grids (Quantifoil Au R1.2/1.3) for cryo-EM data collection. The grids were blotted for 3.5 s and immersed in liquid ethane using Vitrobot (Mark IV, Thermo Fisher Scientific) in condition of 100 % humidity and 8 ℃. The imaging system comprised of Titan Krios operating at 300 kV, Gatan K3 Summit detector, and GIF Quantum energy filter with a 20 eV slit width. Movie stacks were automatically acquired in super-resolution mode (81,000× magnification) using AutoEMation^4^, with a defocus range from -0.5 μm to -3.0 μm. Each stack was exposed for 2.56 s with 0.08 s per frame, resulting in 32 frames and approximately 50 e^-^/Å^2^ of total dose.

**Cryo-EM data processing**

For data processing of TMEM120A, 8168 movie stacks were motion-corrected with 2-fold binning by MotionCor2^5^. Patch-based CTF parameters of the dose-weighted micrographs (1.087 Å per pixel) were determined by cryoSPARC v2^6^, and around three million particles were automatically picked applying Topaz pipeline with a pretrained neural network model (ResNet8, 32 units)^7^. Three rounds of 2D classification enriched images of good classes, resulting in a total of 279,942 particles being selected with box size of 220 pixels. Following an Ab-Initio reconstruction for initial map generation, homogeneous and non-uniform refinement jobs^8^ pushed the resolution to 4.3 Å and 3.8 Å, respectively, under C2 symmetry. To improve map quality, we exported the particles of cryoSPARC refinement jobs, and performed a 3D classification without orientation searching using RELION 3.0 (--skip-align flag)^9^^,^^10^. The best class out of five that exhibits the most intact density with high resolution details, were selected for subsequent homogeneous and non-uniform refinements back to cryoSPARC v2, pushing the resolution to 3.7 Å. Local refinements with masks for TMD and CCD, further improved the local maps at 3.4 Å and 3.8 Å resolution, respectively.

The dataset of TMEM120B was processed in a similar way. Using 2D projections of TMEM120A reconstruction as templates, about 3.7 million particles were autopicked. After 3 rounds of 2D classifications, 560 K particles were selected for subsequent non-uniform refinement with C2 symmetry, using the lowpass filtered TMEM120A map (30 Å) as initial model. We then performed a 3D classification job with 5 classes using RELION 3.0 (--skip-align flag). One class of 138 K particles giving rise to the most intact reconstruction was transformed back to cryoSPARC for further refinement, yielding a final map at 4.0 Å resolution. Map resolutions were determined by the gold-standard Fourier shell correlation (FSC) 0.143 criterion using Phenix.mtriage^11^.

**Model building and refinement**

The structure of human TMEM120A was *de novo* built in Coot^12^. Sequence assignment was guided by bulky residues such as Phe, Tyr and Trp. Most of the side chains were assigned except for peripheral region in the CCD due to limited local resolution. The model of TEME120B was built using a homolog modeling template generated by the structure of TMEM120A. The model was then manually adjusted in Coot.

Subsequently, the models were refined against the corresponding maps by PHENIX^11^ in real space (phenix.real_space_refine) with secondary structure, C2 symmetry and geometry restraints generated by ProSMART^13^. Overfitting of the overall models was monitored by refining the models in one of the two independent half maps from the gold-standard refinement approach and testing the refined model against the other map^14^. Statistics of 3D reconstruction and model refinement can be found in Table S1.

**Supplementary Figs**
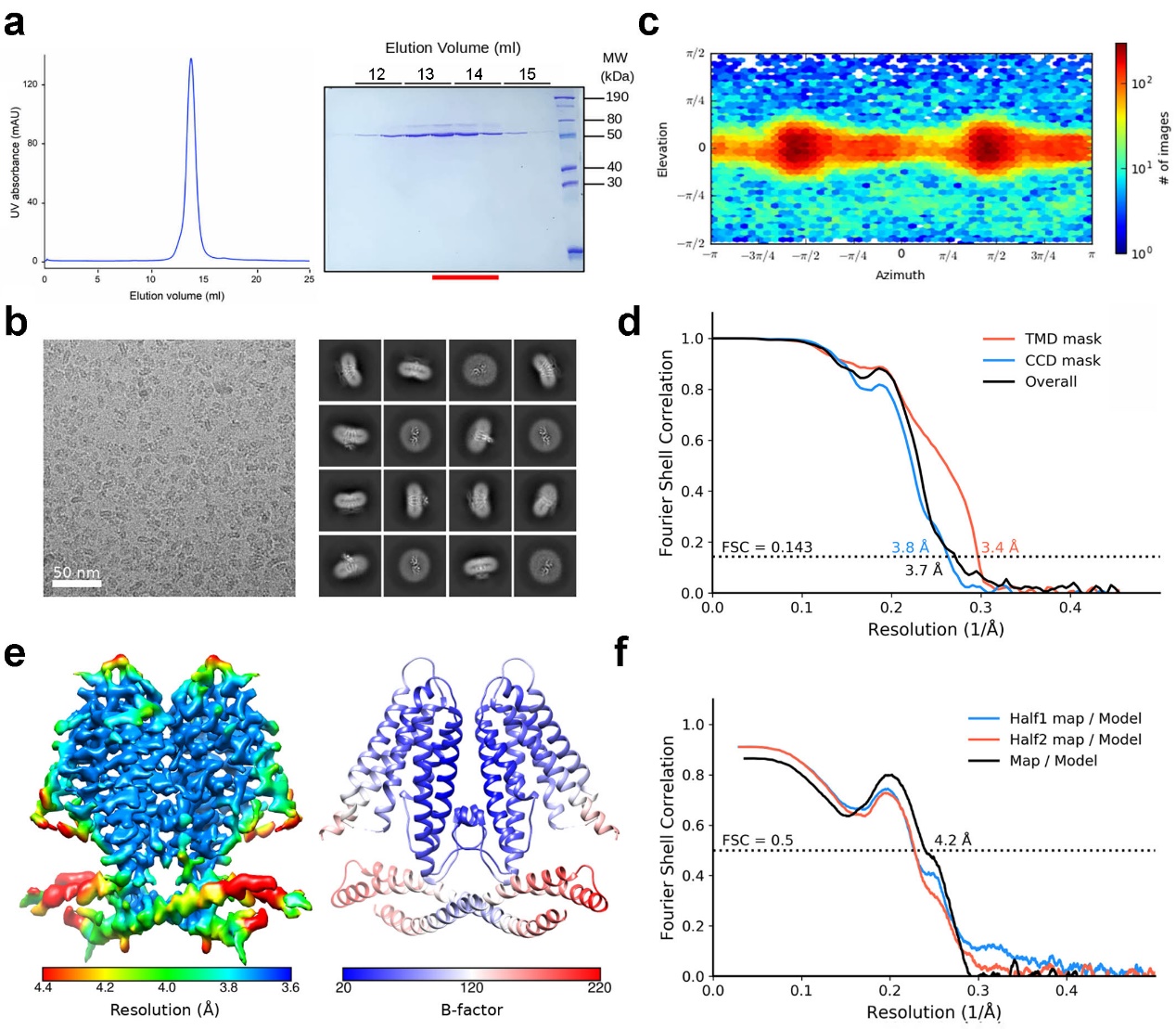

**Fig. S1 | Cryo-EM analysis of human TMEM120A. a,** Last step purification of recombinantly expressed human TMEM120A. A representative chromatogram of gel filtration purification is shown. The indicated fractions were resolved by SDS-polyacrylamide gel electrophoresis followed by Coomassie blue staining. The fractions indicated by red line were collected for cryo-EM analysis. **b,** Representative micrograph and two-dimensional class averages. Scale bar, 50 nm; Box size: 240 Å; Circle mask: 200 Å. **c,** Angular distribution of the particles of the final reconstruction generated by cryoSPARC^6^. **d,** Gold standard Fourier shell correlation (FSC) curves for the 3D reconstructions. The curves for the reconstructions with masks for the entire protein (overall), transmembrane domain (TMD), and coiled-coil domain (CCD) are indicated by black, red, and blue lines, respectively. **e,** Local resolution map estimated by cryoSPARC and generated in Chimera^15^ (*left*) and B-factor heatmap of the structure model against the 3.7 Å reconstruction (*right*). **f,** Validation of the final structure models. FSC curves of the final refined model versus the overall map that it was refined against (black); of the model versus the first half map (blue); and of the model refined in the first half map versus the second half map (red).

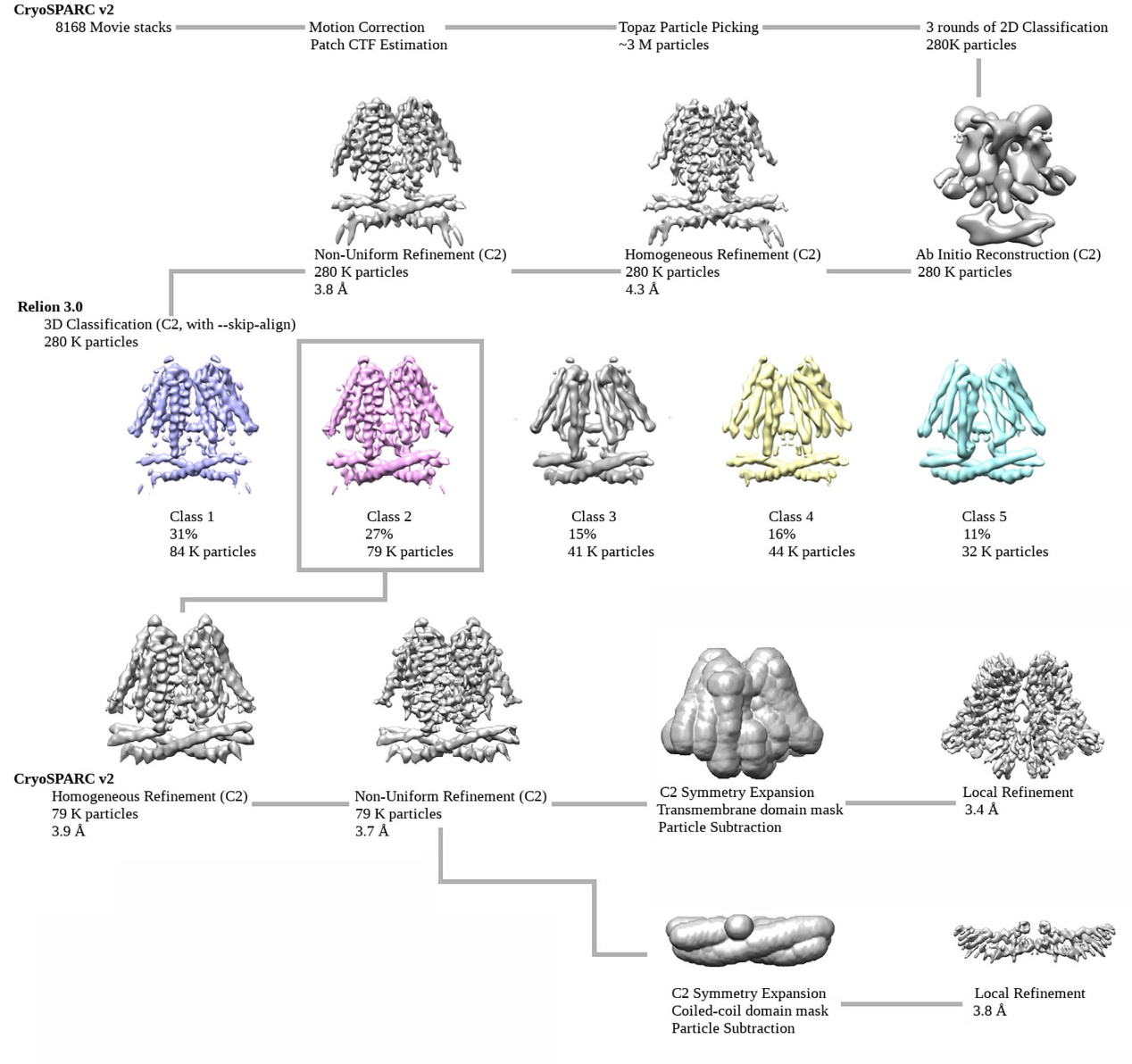

**Fig. S2 | Flowchart for TMEM120A EM data processing.** Please refer to the “Image processing” session in Methods for details.

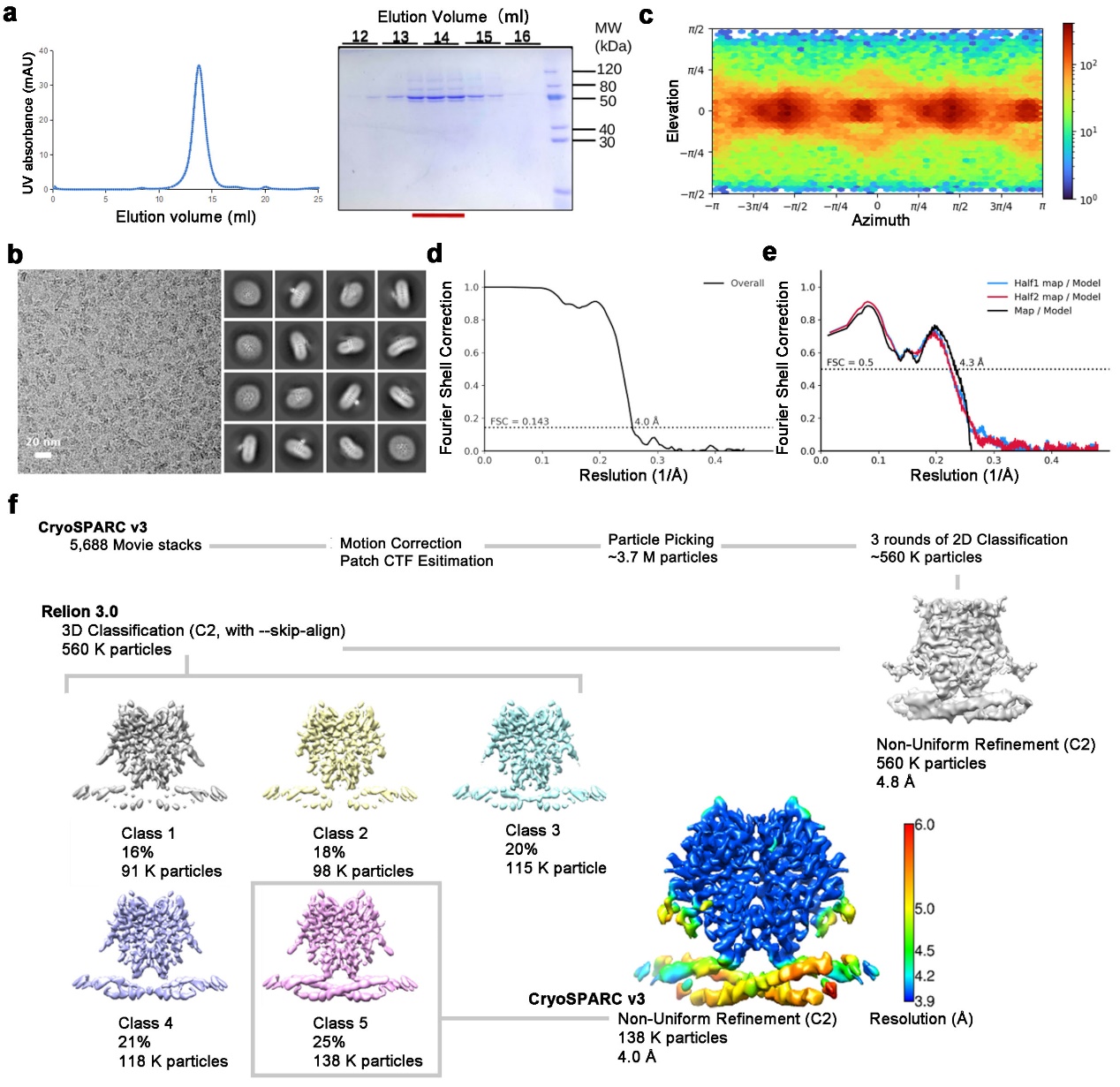

**Fig. S3 | Cryo-EM analysis of human TMEM120B.** **a,** Last step purification of recombinantly expressed human TMEM120B. The chromatogram of gel filtration and corresponding SDS-PAGE of peak fractions stained by Coomassie blue are shown. The fractions highlighted by red line were collected for cryo-EM sample preparation. **b,** Representative micrograph and two-dimensional class averages for TMEM120B. Scale bar, 20 nm; Box size: 240 Å; Circle mask: 200 Å. **c,** Angular distribution of the particles of the final reconstruction generated by cryoSPARC. **d,** FSC curve for the final reconstructed map. **e,** FSC curves for validation of TMEM120B model. **f,** Flowchart for EM data processing of TMEM120B. Local resolution of the final reconstruction is shown. Please refer to the Methods section for details.

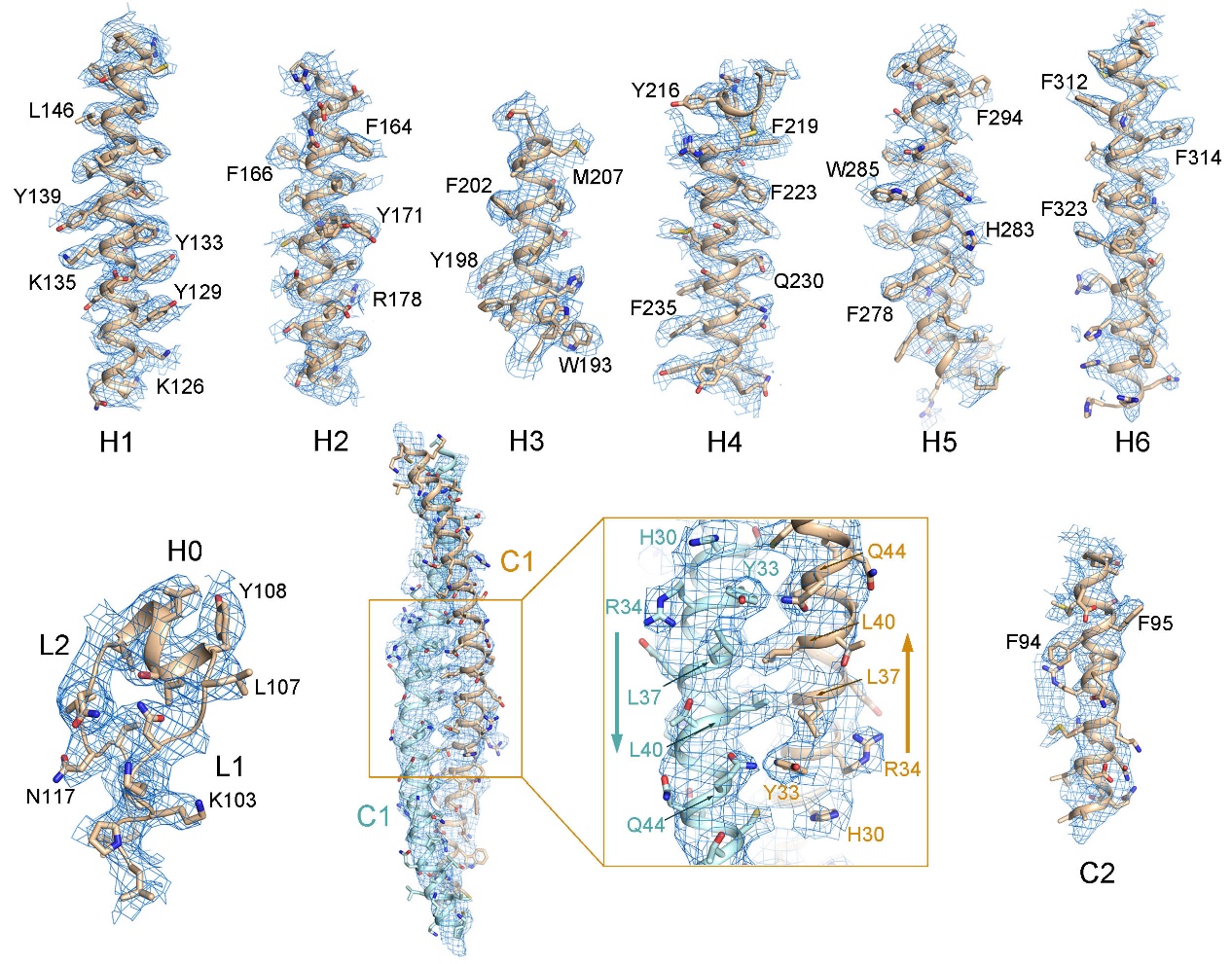

**Fig. S4 | EM maps of human TMEM120A.** Electron density maps for each segment of TMEM120A. The side groups of representative bulky residues are labeled. The densities, shown as blue meshes, are contoured at 4-5 σ in PyMOL^16^.

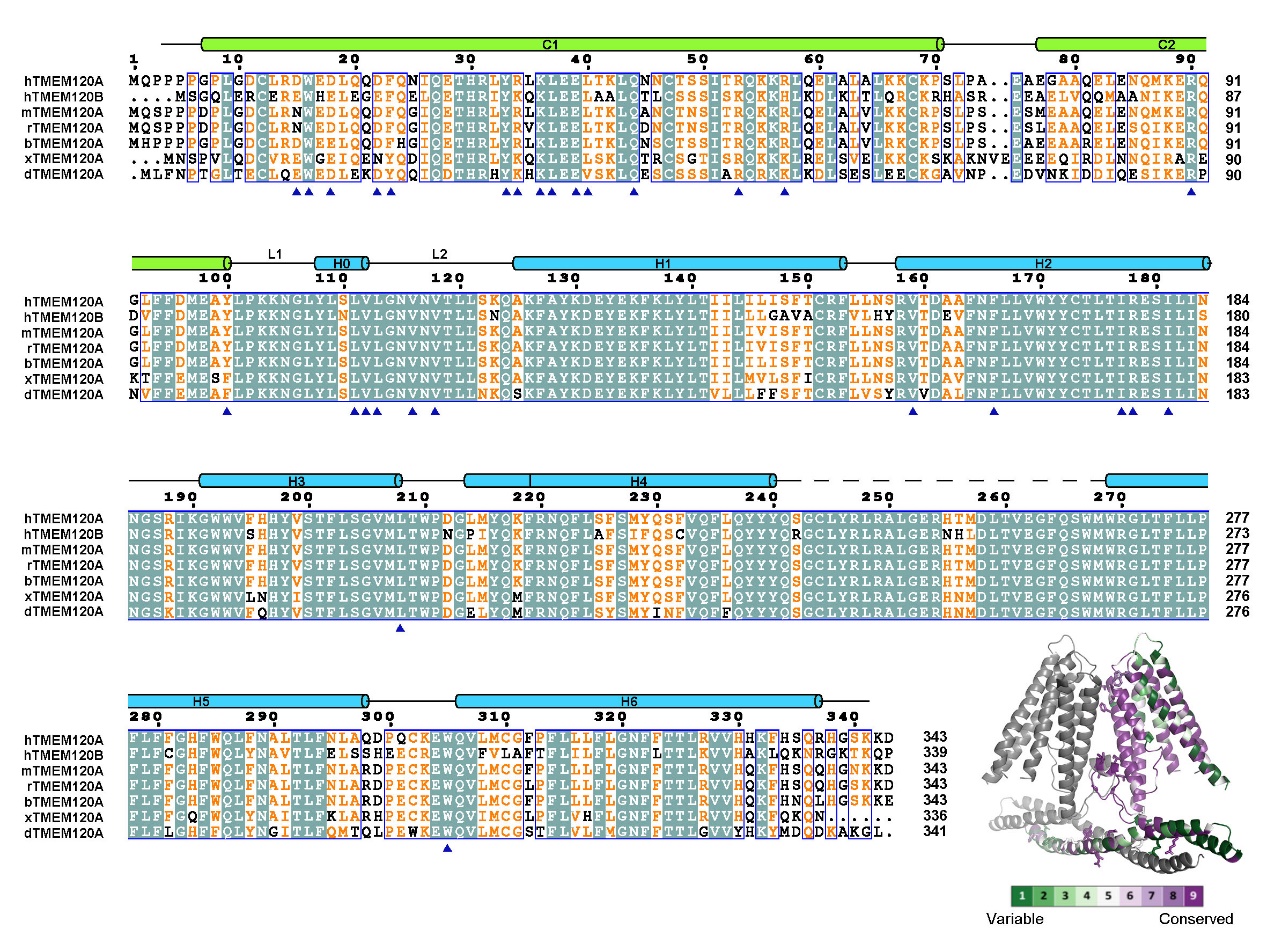

**Fig. S5 | Sequence alignment of TMEM120A from different species and human TMEM120B.** TMEM120A is highly conserved among species. The primary sequences of six TMEM120As from different species and human TMEM120B are aligned using Clustal W^17^. The invariant residues are shaded dark green and the conserved residues are colored orange. Secondary structural elements of human TMEM120A/B are presented above the sequence alignment. The residues that mediate interactions in TMEM120A/B dimer interface are indicated by blue triangles. Based on the alignment, a color scheme of sequence conservation was mapped on a TMEM120A protomer by ConSurf server^18^, as shown in the lower right. Dimer interface residues are shown in sticks. The Uniprot IDs for the aligned sequences are: human TMEM120A: Q9BXJ8; human TMEM120B: A0PK00; mouse TMEM120A: Q8C1E7; rat TMEM120A: Q5HZE2; bovine TMEM120A: Q05B45; frog TMEM120A (xTMEM120A): A1L2R7; zebrafish TMEM120A (dTMEM120A): A3KNK1.

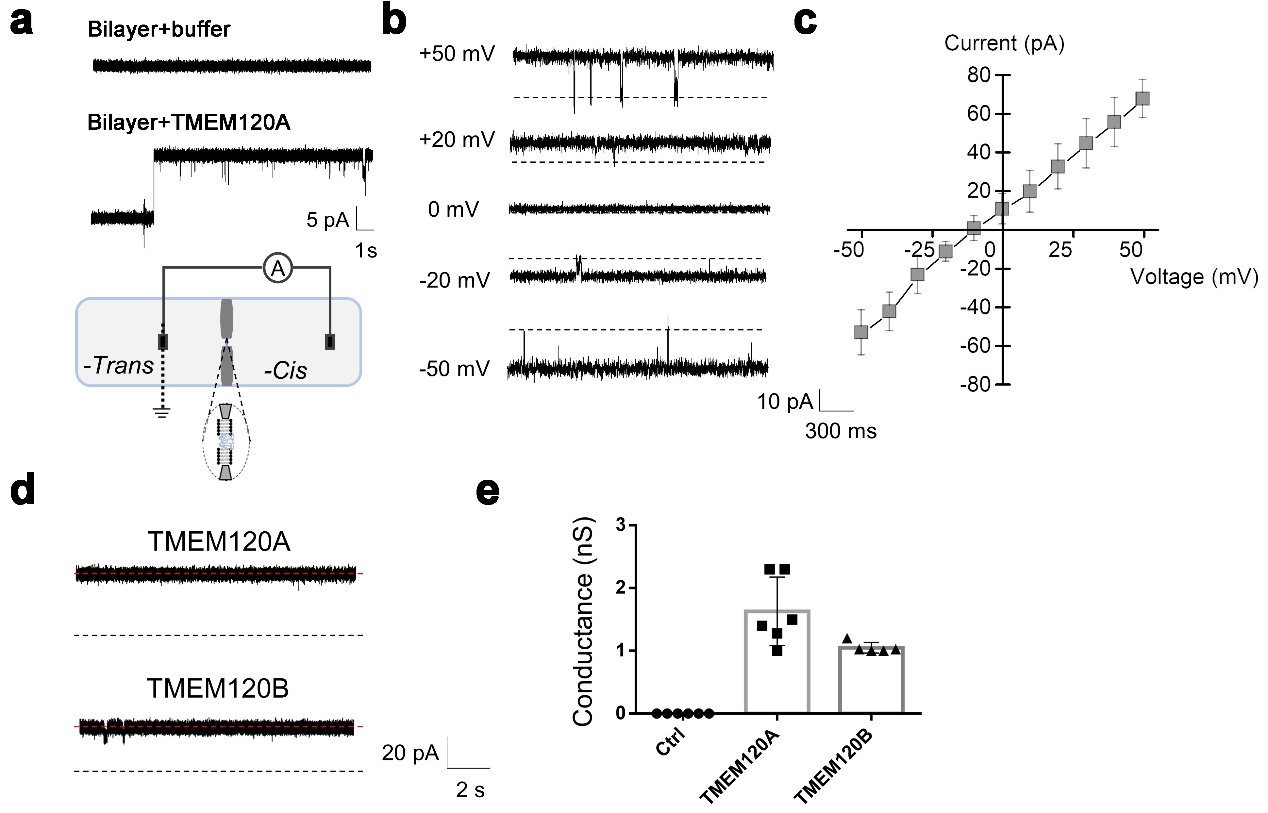

**Fig. S6 | Electrophysiological recordings in lipid bilayer. a,** TMEM120A mediates conducting current in the planar lipid bilayer system. The arrows indicate the point of voltage application at 10 mV. The diagram at the bottom shows the experimental setup for *in vitro* lipid bilayer electrophysiological characterization. **b,** Representative current traces of TMEM120A in response to voltage pulses. **c,** I-V relationship. The I-V curve of TMEM120A is nearly linear (n=3). **d,** Representative current traces of TMEM120A and TMEM120B at 20 mV. **e,** Scatter plot of the ion conductance by TMEM120A and TMEM120B. Control: Protein buffer.

**Table S1 | Statistics for data collection and structural refinement.**

|  | TMEM120A  (PDB: 7CXR;  EMDB-30495) | TMEM120B  (PDB: 7F73;  EMDB-31484) |
| --- | --- | --- |
| **Data collection** |  |  |
| EM equipment | Titan Krios | Titan Krios |
| Voltage (kV) | 300 | 300 |
| Detector | Gatan K3 | Gatan K3 |
| Energy filter | Gatan GIF Quantum,  20 eV slit | Gatan GIF Quantum,  20 eV slit |
| Pixel size (Å) | 1.087 | 1.087 |
| Electron dose (e^-^/Å^2^) | 50 | 50 |
| Defocus range (μm) | -0.5~-3.0 | -0.5~-3.0 |
| Number of images | 8,168 | 5,688 |
| **Reconstruction** |  |  |
| Software | RELION 3.0 / cryoSPARC v2 | RELION 3.0 / cryoSPARC v3 |
| Number of used Particles | 79,080 | 137,998 |
| Symmetry | C2 | C2 |
| Final Resolution (Å) | 3.4-3.7 | 3.9-6.0 |
| Map sharpening B-factor (Å^2^) | -113.7 | -166.2 |
| **Model building and refinement** |  |  |
| Model building software | Coot | Coot |
| Refinement software | Phenix | Phenix |
| **Model composition** |  |  |
| Protein residues | 614 | 586 |
| Side chains | 614 | 586 |
| **Validation** |  |  |
| R.m.s deviations |  |  |
| Bonds length (Å) | 0.007 | 0.005 |
| Bonds Angle (˚) | 0.865 | 1.019 |
| Ramachandran plot statistics (%) | |  |
| Preferred | 93.1 | 97.2 |
| Allowed | 6.9 | 2.8 |
| Outlier | 0.0 | 0.0 |
